# Down-regulated Long Noncoding RNA *HOXA11-AS* affects trophoblast cell proliferation and migration by regulating *RND3* and *HOXA7* expression in preeclampsia

**DOI:** 10.1101/285643

**Authors:** Yetao Xu, Dan Wu, Jie Liu, Zhonghua Ma, Bingqing Hui, Jing Wang, Yanzi chen, Sailan Wang, Yifan Lian, Lizhou Sun

**Author notes:** **Corresponding author:** To whom correspondence should be addressed to Lizhou Sun. These authors contributed equally to this work and should be regarded as joint first authors (Yetao Xu, Dan Wu, Jie Liu).

## Abstract

The long noncoding RNA *HOXA11-AS* reveals abnormal expression in numerous human diseases. However, its function and biological mechanisms remain unclear in Preeclampsia (PE). In this study, we report that *HOXA11-AS* was significantly downregulated in preeclampsic placental tissues and could contribute to the occurrence and development of Preeclampsia. Silencing of *HOXA11-AS* expression could significantly suppress trophoblast cell growth and migration, whereas *HOXA11-AS* overexpression facilitated cell growth in HTR-8/SVneo, JEG3 and JAR cell lines. RNA-seq analysis also indicated that *HOXA11-AS* silencing preferentially regulated numerous genes associated with cell proliferation and cell migration. Mechanistic analyses showed that *HOXA11-AS* could recruit Ezh2 and Lsd1 protein, and regulate *RND3* mRNA expression in nucleus. In cytoplasm, *HOXA11-AS* modulate *HOXA7* expression by sponged miR-15b-5p, thus affecting trophoblast cell proliferation. Together, these resulting data confirm that aberrant expression of *HOXA11-AS* is involved in the occurrence and development of Preeclampsia, and may act as a prospective diagnosis and therapeutic target in PE.

## Introduction

Preeclampsia (PE), characterized by blood pressure higher than 140/90 mmHg after 20 weeks in pregnancy, is a major contributor of pregnancy-related-death and fetal morbidity. PE has been afflicting nearly 3-5% of pregnancies, especially in developing countries [1, 2]. Despite the considerable morbidity and mortality, the cause of preeclampsia has been a mystery. Delivery of the placenta is the only known remedy for PE [3], while other effective prevention strategies have not yet been developed. The preferential clinical treatment against PE is to use combination of labetalol with magnesium sulfate to slow down the progression of this disorder and expectantly extend the pregnancy period. With in-depth study, PE, which results from the aberrant expression of numerous PE-associated genes [4-7], could be considered as a heterogeneous disease with diverse clinical and molecular characteristics. Therefore, a deepening understanding of the biological mechanisms on PE might furnish more preferences for diagnose and treatment.

LncRNAs, which is longer than 200-bp with little or no protein-coding capacity, have intrinsic function as RNA [8, 9]. Recently, technological advances have allowed the analysis of long noncoding RNAs (lncRNAs) in diverse human diseases. Emerging studies have demonstrated that lncRNAs have been implicated in a variety of biological and pathological processes, including cell differentiation [10], cell metabolism[11], immune response[12] and disease-associated development [13-15]. Additionally, the aberrant levels of lncRNAs have been reported, which have positively or negatively affected various gene expressions in diverse human diseases, including PE[16-19]. Furthermore, lots of studies demonstrated that lncRNAs could act as significant regulatory molecules to regulate related-gene expression at different levels, such as chromatin modification, transcriptional and post-transcriptional modification [9, 20]. For instance, lncRNA *CCAT1* modulate *SPRY4* and *HOXB13* expression by binding to *SUV39H1*(suppressor of variegation 3-9 homolog 1) and *EZH2* (enhancer of zeste 2 polycomb repressive complex 2 subunit) to affect cell growth and migration in esophageal squamous cell carcinoma[21]. Apart from their role in gene expression regulation, lncRNAs could also crosstalk with associated-gene expression by competing for shared microRNAs in post-transcriptional levels to affect the occurrence and development of various diseases [22-24].

*HOXA11-AS*, a 1628-bp lncRNA gene located on chromosome 7p15.2, plays significant roles in various disorders[25-28]. For instance, *HOXA11-AS* could promote cell growth and invasion of gastric cancer through interacting with *EZH2* and *LSD1* (Histone demethylase lysine specific demethylase 1)[29]. In addition, *HOXA11-AS* could compete for shared miR-140-5p to promote the Glioma tumorigenesis [30]. However, the biological functions of *HOXA11-AS* in PE remains unclear, which impels us to further explore the role and molecular mechanism of *HOXA11-AS* in PE.

In this study, we demonstrated that the expression level of *HOXA11-AS* was significantly downregulated in preeclampsic placental tissues compared with that in normal tissues. Furthermore, knockdown of *HOXA11-AS* could impair cell growth and migration in various trophoblast cell lines. Associated mechanistic exploration demonstrated that *HOXA11-AS* could exhibit different regulatory mechanisms in regulation of *RND3* and *HOXA7* expression in nucleus and cytoplasm, thus involving in the occurrence and development of PE. The unraveling role of HOXA11-AS will provide novel insights for future PE treatments.

## Materials and Methods

### Tissue samples and Ethics statement

60 PE patients were selected in this study at the obstetrical department of First Affiliated Hospital of Nanjing Medical University. All patients signed written informed consent. The clinical characteristics of the PE patients were collected in Table1. This research was authorized by the Ethnics Board of the First Affiliated Hospital of Nanjing Medical University, China, and it was performed in compliance with the Declaration of Helsinki Principles.

### Cell culture

In this study, HTR-8, JEG3 and JAR cell lines were purchased from the Institute of the Chinese Academy of Sciences (Shanghai, China). HTR-8 and JAR were cultured in 1640 (GIBCO, Nanjing, China) medium supplemented with 10% fetal bovine serum (FBS) (GIBCO, BRL, Invitrogen, Carisbad, CA, USA), 100U/ml penicillin and 100 mg/ml streptomycin. JEG3 was cultured in MEM with 10% FBS. All cell lines were cultured in humidified air at 37 **°**C/5% CO_**2**_.

### Plasmid construction

Full-length complementary DNA of *HOXA11-AS* (1628bp, NR_002795.2), *RND3* sequence (put base pairs and NR number here) were synthesized and cloned into the pcDNA3.1(+) plasmid vector (Invitrogen). Resulting plasmids as well as empty pcDNA3.1(+) vector were transfected into HTR-8/SVneo, JEG3 and JAR cells, respectively on 6-well plates and/or 24-well plates.

### Cell transfection

Lipofectamine 2000 or/and Lipofectamine 3000 transfection reagent (Life Technologies, Invitrogen, USA) were used to transfect the trophoblast cell lines with siRNAs targeting *HOXA11-AS* mRNA according to the manufacturer’s protocol. All the siRNA sequences were listed in Supplementary Table 1. Interference target sequences of *HOXA7* and *RND3* were purchased from Invitrogen. The transfected cells on 6-well plates were harvested for further experiments 48 h post-transfections.

**Table 1:** Clinical characteristics of preeclamptic and normal pregnancies.

| Variable | PE (N=60) | Normal (N=60) | *P* value<sup>a</sup> Normal vs P |
| --- | --- | --- | --- |
| Maternal age (year) | 32.23±4.526 | 34.62±3.3763 | p>0.05 |
| Maternal weight (kg) | 75.99±10.995 | 74.467±7.979 | p>0.05 |
| Smoking | 0 | 0 | p>0.05 |
| Systolic blood pressure (mm Hg) | 162.83±15.103 | 116.6±7.983 | p<0.01 |
| Diastolic blood pressure (mm Hg) | 106.417±10.807 | 71.54±8.767 | p<0.01 |
| Proteinuria (g/day) | >0.3g | <0.3g | p<0.05 |
| Body weight of infant (g) | 2287.33±908.078 | 3385.42±372.269 | p<0.05 |
| Gestational age (week) | 33.93±3.231 | 38.38±0.958 | p<0.05 |

### RNA extraction and qRT-PCR analyses

Total RNA from each treatment were extracted by using the TRIzol reagent (Thermo Fisher Scientific) and qRT-PCR analyses were conducted by using the SYBR® Green Master Mix (TaKaRa BIO INC, Otsu, Japan) as protocol described. The sequences of specific primers used are shown in Supplementary Table 1.

### Subcellular fractionation location

The nuclear and cytosolic fractions were separated and purified using the PARIS Kit (Life Technologies, Carlsbad, CA, USA) according to manufacturer’s manual.

### Cell viability assays

Cell viability was detected with MTT kit (Sigma) following the manual. For colony formation assay, 600,/800,/1000 cells treated with siRNAs and/or plasmid were plated on 6-well plates and maintained in proper media containing 10% FBS for 10-14 days, during which the medium was replaced every 4 days. Colonies were then fixed with methanol and stained with 0.1% crystal violet (Sigma) in PBS for 30 min. Colony formation was determined by counting the number of stained colonies.

BrdU experiments were performed by using a BrdU Cell Proliferation Assay Kit (Millipore, Cat.No.2750) following the protocol. The higher OD reading represents the higher BrdU concentration in each sample.

EdU assay was implemented as a complementary method in order to authenticate the proliferation level. Usually, we exploit the 5-ethynyl-2-deoxyuridine labeling/detection kit (Ribobio, Guangzhou, China) to evaluate cell proliferation, following the manufacturer’s manual.

### Flow-cytometric analysis of cell cycle and apoptosis

After transfecting cells with siRNAs or plasmid, we performed FITC-Annexin V and Propidium iodide (PI) staining by using the FITC-Annexin V Apoptosis Detection Kit (BD Biosciences, Franklin Lakes, NJ, USA) following manufacter’s instruction. Cell cycle level was determined by propidium oxide staining using the Cycle TEST PLUS DNA Reagent Kit (BD Biosciences, Franklin Lakes, NJ, USA) following the protocol and analyzed by FACScan. The ratio of the cells in G0/G1, S and G2/M phase were calculated and compared.

### Transwell assays

The ability of cell migration and invasion was detected and analyzed by transwell assays as previously reported in Xu et al[31]. 24-well chambers were placed into the upper chamber of an insert with 8µm pore size polycarbonate membrane (Millipore, Billerica, MA, USA). Medium containing 15% FBS was added to the lower chamber. After incubation for 24-48 h, the cells on the upper membrane were removed with cotton swab. Cells that migrated or invaded through the polycarbonate membrane were stained with methanol and 0.1% crystal violet. Experiments were conducted repeatedly three times.

### Western blot assays

WB assays were conducted as previously reported in Xu et al[31], and the following primary antibodies were used respectively: Anti-EZH2, Anti-LSD1, Anti-AGO2 (Millipore, USA), Anti-RND3 and Anti-HOXA7 from Proteintech (WuHan, China).GAPDH (Cell Signaling, San Jose, CA, USA) antibody was used as the control.

### RNA-seqbioinformatic analysis

The RNA-seq experiments were conducted by Wuhan Genomics Institute (Wuhan, China). Culture cells which were treated were extracted. To stablish the mRNA-seq library, the cDNAs were fragmented via nebulization following the protocol.

### Chromatin immunoprecipitation assays (ChIP)

Chromatin immunoprecipitation (ChIP) assays were conducted as previously described in Xu et al [31]. According to the manual, experiments were performed by using EZ-CHIP KIT (Millipore, USA). Relevant antibodies against H3K27me3, H3K4me2 and other target proteins were purchased from Millipore. The primer sequences of gene promoter were summarized in Supplementary Table 1. Quantification of immunoprecipitated DNA were detected and analyzed by qRT-PCR. Experiments were repeated three times.

### RNA immunoprecipitation assays (RIP)

RNA immunoprecipitation (RIP) experiments were performed following the manufacturer’s protocol of Magna RIP ™ RNA-Binding Protein Immunoprecipitation Kit (Millipore, USA). Antibodies, including EZH2, SUZ12, DNMT3a, DNMT3b, LSD1, AGO2 and STAU1, were purchased from Millipore.

### Luciferase reporter assays

Luciferase reporter assays were performed as previously reported in Zhang et al[21]. HOXA11-AS and HOXA7 3′-UTR cDNA fragments were amplified by PCR assays, and then subcloned downstream of the luciferase genes in the pGL3 plasmid. Mutant of plasmids, such as pGL3-HOXA7-3 ′ UTR MUT and pGL3-HOXA11-AS-MUT, were obtained by platinum pfx DNA polymerase according to the protocol. Luciferase activity was measured by the Dual-Luciferase Reporter Assay System (Promega, Madison, WI, USA). Briefly, 1×10**5** HTR-8/SVneo cells were plated in 24-well plates for 36h. After 48 hours post-transfection, the cells were obtained and lysed for further experiments. The relative luciferase activity was normalized with renilla luciferase activity.

### Statistical analysis

All statistical analyses in our experiment were performed by using SPSS 20.0 software (IBM, SPSS, USA). These resulting data was represented as the mean **±** SD. Statistical significance were ascribed at P<0.05(*) or P<0.01(**). Each experiment was repeated at least three times independently.

## Result

### *HOXA11-AS* is downregulated in human preeclampsic tissues

The expression level of *HOXA11-AS* was analyzed in 60 preeclamptic tissues and normal tissue samples by qRT-PCR. We found that the *HOXA11-AS* expression was significantly downregulated in preeclampsic tissues (Figure1A). The detailed clinical characteristics of the patients who meet criteria were listed in table 1. In addition, we discovered that there were no significant differences between preeclampsia and the normal in gestational age and maternal age. (P>0.05). On the contrary, there were significant differences in systolic blood pressure, diastolic blood pressure and body weight of infant between preeclampsia and the normal (P<0.05).

### *HOXA11-AS* regulates trophoblast cell proliferation and migration *in vitro*

Since human lncRNAs play essential roles in various cellular processes, we detected the expression of *HOXA11-AS* in four trophoblast cell lines and another two cell lines related to pregnancy, including HTR-8/SVneo, BeWo, JEG-3 and JAR, WISH and HUVEC-C. As shown in figure 1B, we found that the relative *HOXA11-AS* level in HTR-8/SVneo cells was higher than that in other cell lines, whereas the expression levels of *HOXA11-AS* in BeWo, JEG3 and JAR cell lines were relatively lower compared to those in WISH and HUVEC-C cell lines.

**Figure 1:**
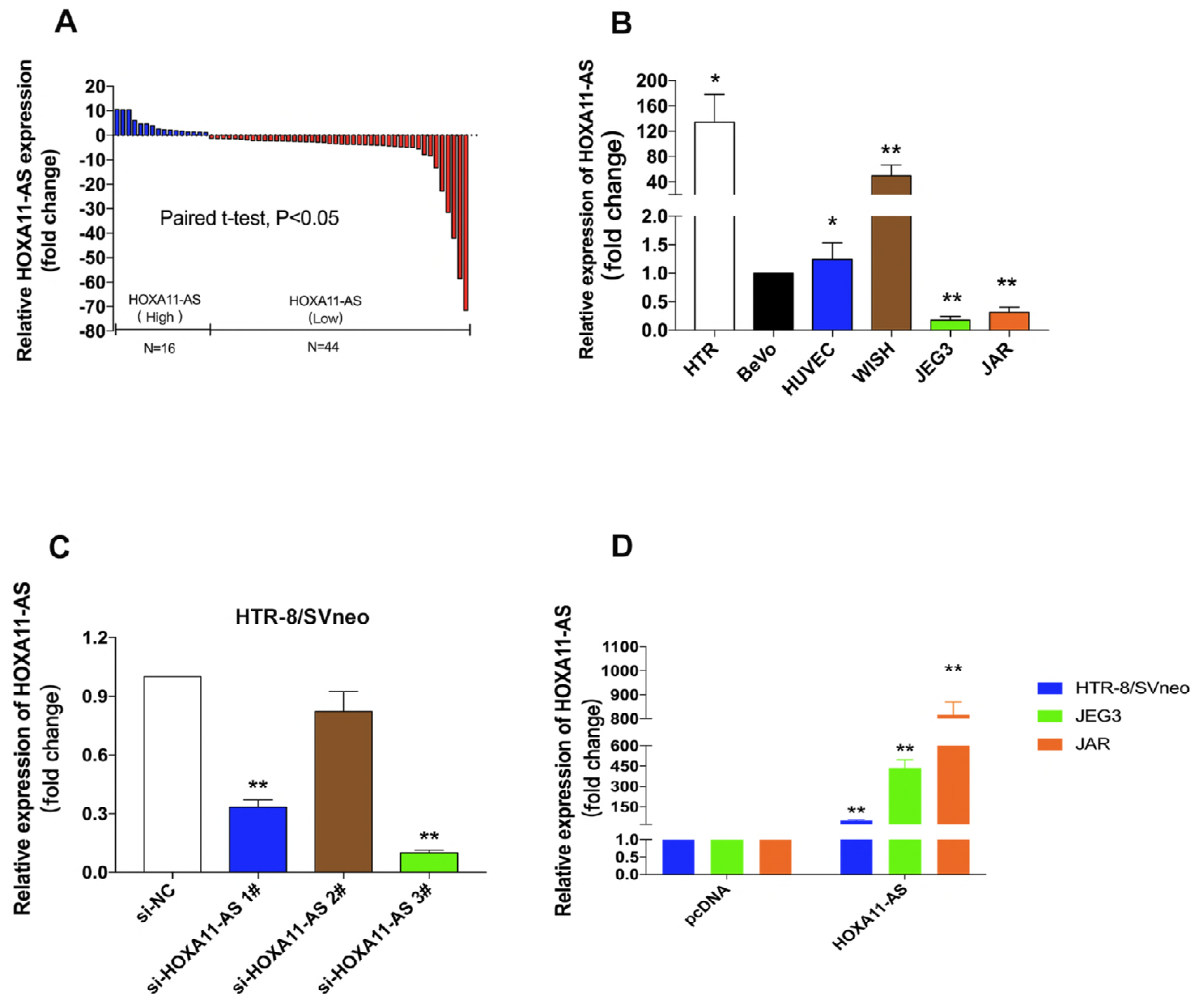
Relative *HOXA11-AS* expression in PE. (A) The relative expression of *HOXA11-AS* was measured by qRT-PCR. The levels of *HOXA11-AS* were lower in Preeclamptic placentas samples (N=60) than that in normal placentas (N=60). (B) *HOXA11-AS* expression were detected by qRT-PCR in several cell-lines and were normalized to that in HTR-8/SVneo. (C and D) The relative expression of *HOXA11-AS* following the treatment of HTR-8/SVneo, JEG3 and JAR cells with siRNAs and pcDNA3.1**+***HOXA11-AS*. At least three times of biological replicates have been performed and presented. (**: P < 0.01, *: P<0.05).

To explore potential role of *HOXA11-AS* in trophoblast cells, we performed overexpression and knowdown model of HOXA11-AS *in vitro*. It was found that the expression levels of *HOXA11-AS* were exogenously influenced by specific siRNAs and overexpression plasmid in HTR-8/SVneo, JEG3 and JAR cell lines (figure 1C and 1D). Then we performed MTT and Colony formation assays to illustrate the effect of *HOXA11-AS* on the proliferation of HTR-8/SVneo, JEG3 and JAR trophoblast cells. The resulting data revealed that silencing of *HOXA11-AS* significantly retarded cell growth compared with that of the controls, while upregulation of *HOXA11-AS* could enhance cell proliferation (Figure 2A and 2B). In addition, Ethynyl deoxyuridine (EdU) staining assays and BrdU assays also demonstrated that *HOXA11-AS* knockdown inhibited trophoblast cells proliferation; however, *HOXA11-AS* overexpression boosted the rate of proliferating trophoblast cells (Figure 2C and 2D). These data indicate that down-regulated *HOXA11-AS* might play the role as a suppressor in the inhibition of trophoblast cell proliferation.

**Figure 2:**
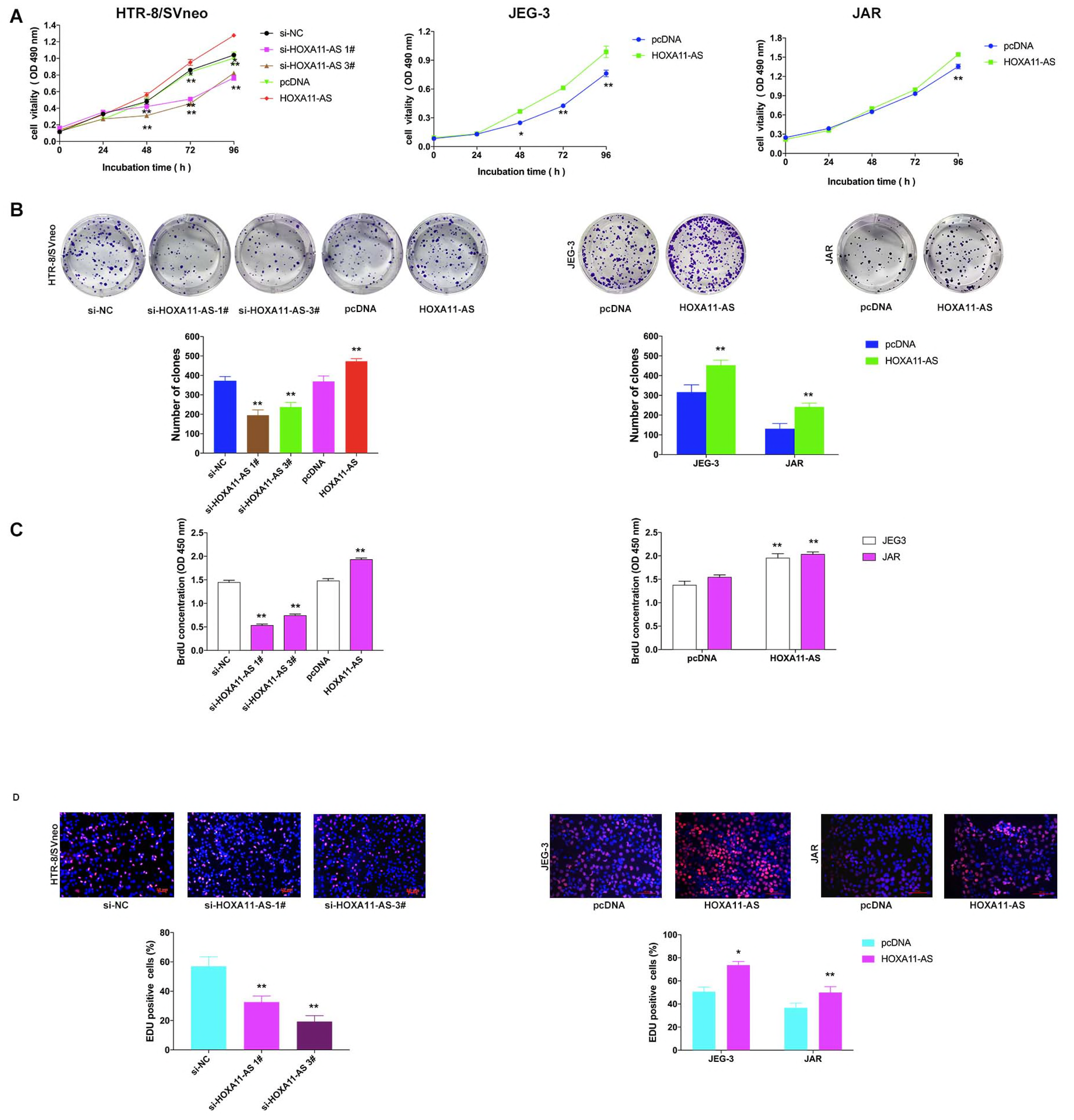
Effect of *HOXA11-AS* on proliferation in trophoblast cells. (A) MTT assays were used to determine the viability of si-*HOXA11-AS*-transfected trophoblast Cells. (B) Colony formation assays were performed to determine the proliferation of si-*HOXA11-AS*-transfected HTR-8/SVneo, JEG-3 and JAR cell lines. Colonies were counted and captured. (C) BrdU assays were used to detect the cell proliferation after transfection, respectively. (D). Proliferating Trophoblast Cells were labeled with EdU. The Click-it reaction revealed Edu staining (red). Cell nuclei were stained with DAPI (blue). (**: P < 0.01, *: P<0.05)

Furthermore, transwell assays confirmed that silencing of *HOXA11-AS* significantly inhibited trophoblast cell migration and invasion compared with the si-NC treatment (Figure 3A). In contrast, upregulated *HOXA11-AS* could stimulate cell migration and invasion (Figure 3A).

**Figure 3:**
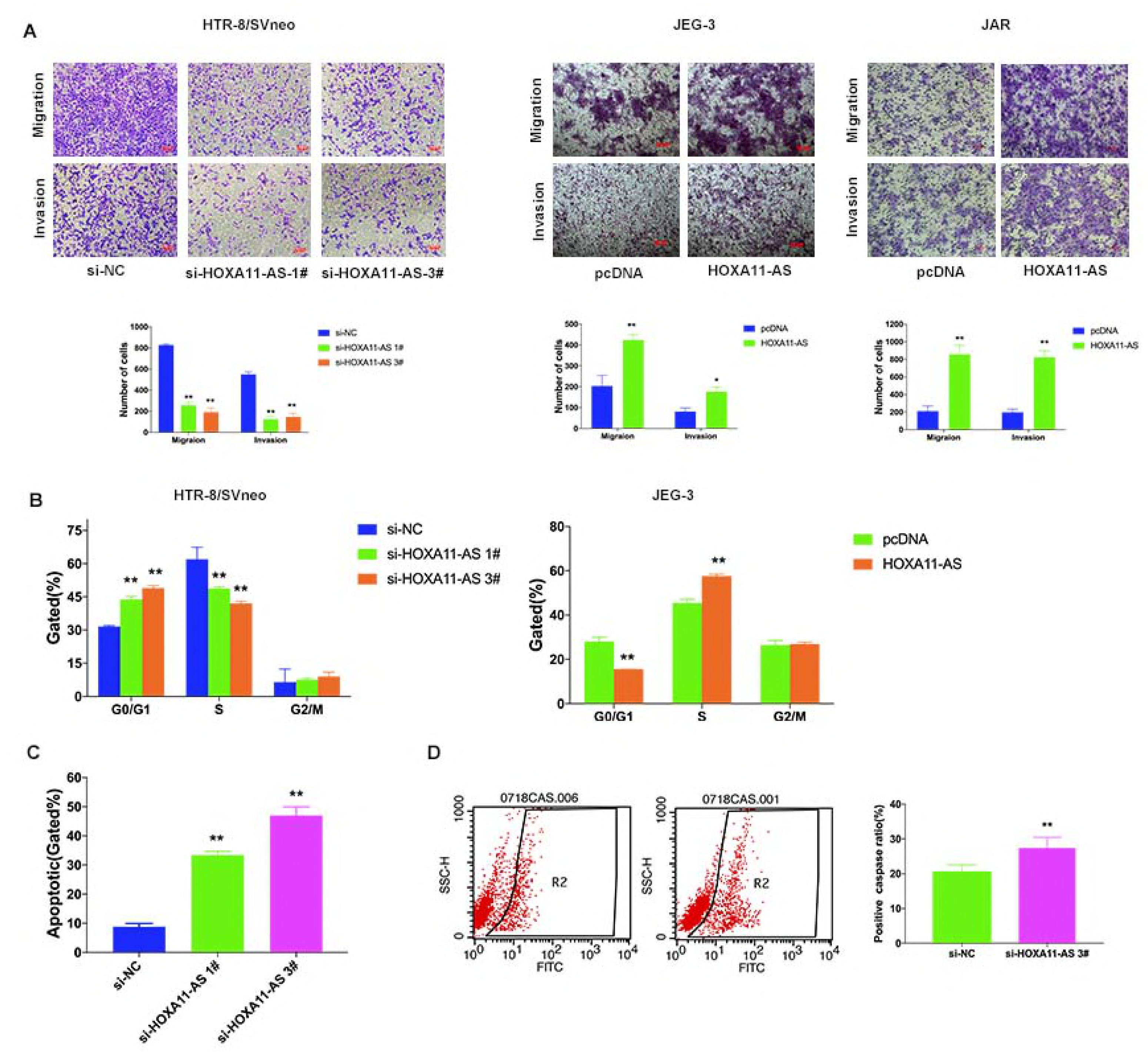
*HOXA11-AS* regulates trophoblast cell migration, cell cycle and apoptosis in vitro. Trophoblast Cells were treated with specific *HOXA11-AS* siRNAs and/or overexpression plasmid (A) Transwell assays were used to investigate the changes in migratory and invasion abilities of trophoblast cells after transfection, respectively. (B) Cell cycle analyses by Flow cytometry in HTR-8/SVneo and JEG-3 cells. (C) Flow cytometry was used to detect the apoptotic rates of cells. LR, early apoptotic cells; UR, terminal apoptotic cells. (D) Flow cytometry assays to detect the protein expression level of cleaved-caspase after transfected with siRNAs against *HOXA11-AS* in HTR-8/SVneo. All experiments were performed in biological triplicates with three technical replicates. (**: P < 0.01, *: P<0.05, n.s.: not significant)

### Effect of HOXA11-AS on cycle and apoptosis in vitro

Because cell proliferation assays cannot thoroughly reflect cell cycle changes, we next performed flow cytometry analysis to detect whether the cell cycle progression was affected after *HOXA11-AS* knockdown. The results revealed that the cells transfected with specific siRNAs promoted cell accumulation in the G0-G1 phase compared with that treated with si-NC. In contrast, *HOXA11-AS* overexpression could reduce cell cycle accumulation in G0-G1 phase (Figure 3B).

Also, flow cytometry assays were performed to investigate whether knockdown of *HOXA11-AS* affected the cell apoptosis. The results showed that the ratio of total apoptotic cells were dramatically increased in cells transfected with siRNAs in the HTR-8/SVneo (Figure 3C). Similarly, we conducted flow cytometry assays to detect the protein expression level of cleaved-caspase, which further confirmed that the apoptosis level was increased after HOXA11-AS knowdown (Figure 3D). These findings suggest that *HOXA11-AS* promotes proliferation and inhibits apoptosis in trophoblast cells.

### Gene expression profiling

To investigate the *HOXA11-AS*-associated pathway on an unbiased basis in Preeclampsia, we conducted RNA-seq and evaluated the gene expression profiles of HTR-8/SVneo cells transfected with siRNAs against *HOXA11-AS*. After knockdown of HOXA11-AS, 131 mRNAs showed at least a 2-fold increased abundance, whereas a total of (≤ 2-fold) of 99 genes showed decreased abundance (Figure 4A). GO analysis showed many significant biological processes were involved in cell proliferation, migration, as well as apoptosis (Figure 4B). Among all the enriched genes, there are many famous proliferation-related and migration-associated genes, such as *TNFSF9, TFPI2, CA9, IFITM1, TMEM158, RND3, ESM1, NAMPT1, HOXA7, PSAT1, CPA4, MEST, OLR1*, etc. We suspected that some altered genes might induce the occurrence and development of Preeclampsia. The expression changes of these genes, therefore, were selectively demonstrated by qRT-PCR in *HOXA11-AS* depleted and/or *HOXA11-AS* overexpressed HTR-8/SVneo cells and *HOXA11-AS* overexpressed JAR cells (Figure 4C and 4D). Lastly, *RND3* and *HOXA7* have been identified as candidate factors involved in cell proliferation, apoptosis, and migration; therefore, we selected *RND3* and *HOXA7* for further study.

**Figure 4:**
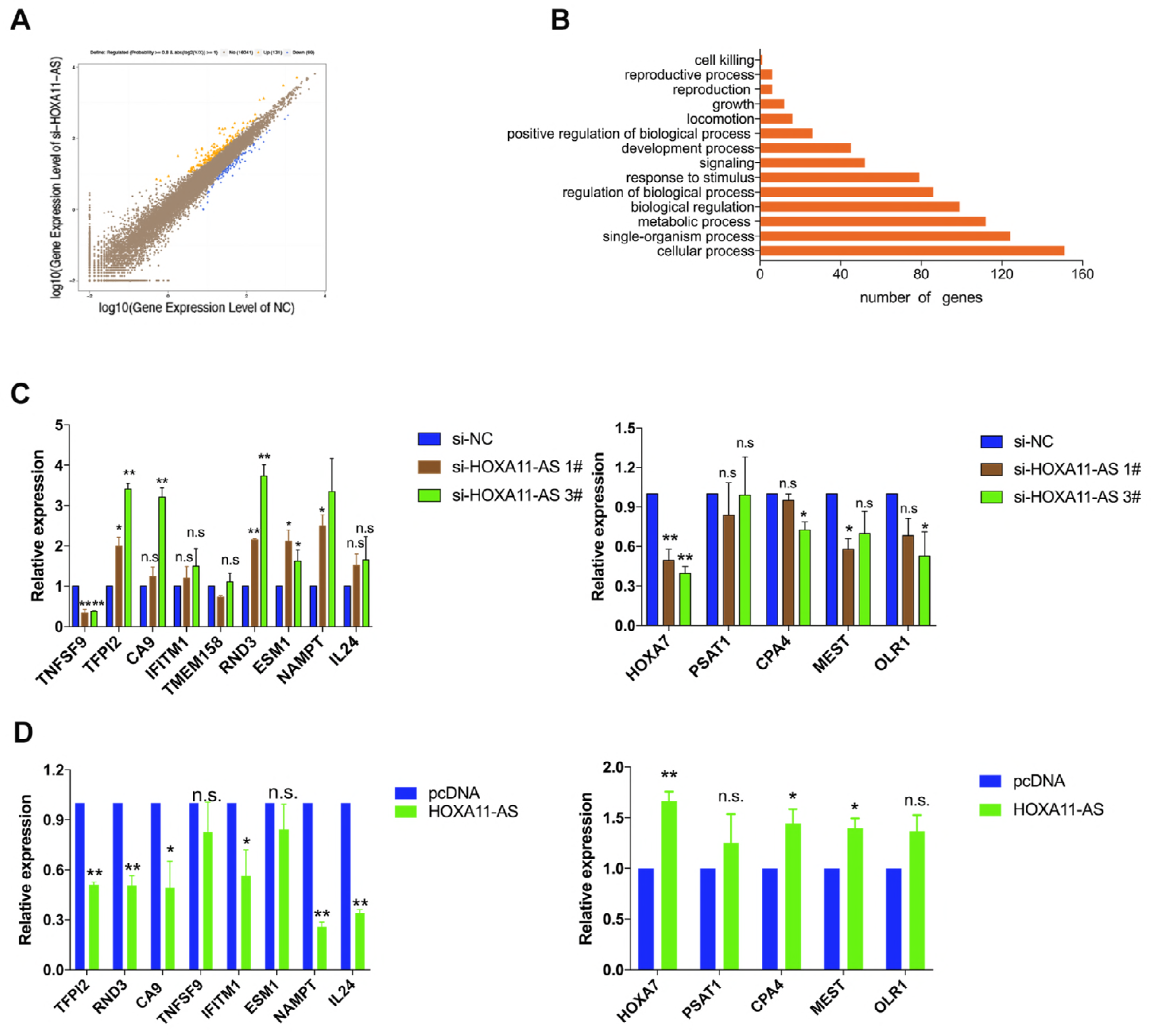
*HOXA11-AS* knockdown increases the expression of genes involved in cell proliferation and migration. (A) RNA transcriptome sequencing analysis was performed to analyze gene expression profiling in HTR-8/SVneo cells following *HOXA11-AS* knockdown. The picture showed the all of different expressed gene. (B) GO analysis for all genes with altered expressions between the scrambled siRNA-treated and si-*HOXA11-AS*-treated cells in vitro. Cell growth was among the significant biological processes for genes whose transcripts level were changed in the *HOXA11-AS*-depleted Trophoblast cells. (C and D) qRT-PCR analysis in si-*HOXA11-AS*-treated trophoblast cells reveal altered mRNA level of genes involved in cell proliferation and migration upon *HOXA11-AS* depletion. (**: P<0.01, *: P<0.05, n.s.: not significant).

### *HOXA11-AS* could recruit Lsd1 and Ezh2 in nucleus, thus epigenetically silencing of *RND3*

To further explore the potential biological mechanisms of *HOXA11-AS* medicated regulation in trophoblast cells, we first performed subcellular fractionation assays to assess the distribution of *HOXA11-AS* in nuclear and cytoplasmic fractions in HTR/SVneo, JEG-3 and JAR cells. As shown in figure 5A, approximately 70% of *HOXA11-AS* is located in the trophoblast nucleus, and 30% of *HOXA11-AS* is in the cytoplasm. Therefore, these findings indicated that *HOXA11-AS* might play an essential role in transcriptional regulation. Then, we employed bioinformatics analysis to predict possibilities of RNA binding proteins, including Ezh2 (H3K27me3), Suz12 (H3K27me3), Lsd1 (H3K4me2), Dnmt1, Hur, Stau1 and Ago2 (http://pridb.gdcb.iastate.edu/RPISeq/references.php) [32]. As shown in Figure 5B, *HOXA11-AS* may interact with Ago2, Ezh2 (H3K27me3) and Lsd1(H3K4me2) in trophoblast cells. And previous studies have reported that *HOXA11-AS* could recruit Ezh2 and Lsd1 to epigenetically silenced targets in many tumor cell lines

**Figure 5:**
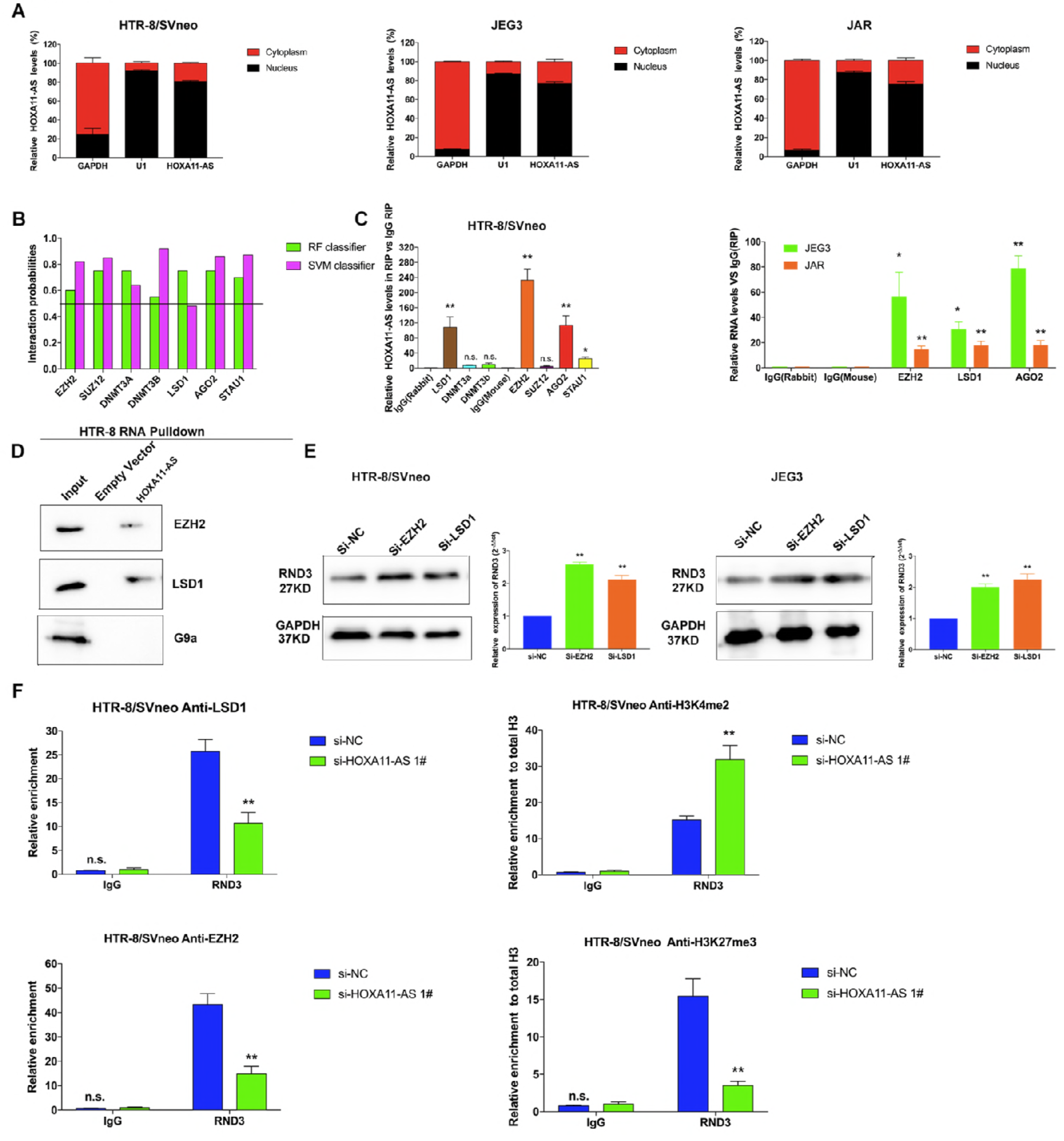
*HOXA11-AS* can recruit *EZH2* and *LSD1* to silence *RND3* expression. (A) Cell fractionation assays indicated that *HOXA11-AS* is mostly located in nucleus. GAPDH and U1 acted as the marker of cytoplasm and nucleus, respectively. (B) Bioinformatics were used to predict this possibility of interaction of *HOXA11-AS*. Predictions with probabilities>0.5 were considered positive. RPISeq predictions are based on Random Forest (RF) or Support Vector Machine (SVM). (C) RIPs experiments were performed and the coprecipitated RNA was detected by qRT-PCR. (D) In vitro transcribed, pull down assays showed that desthiobiotinylation-*HOXA11-AS* could retrieve *EZH2* and *LSD1* in HTR-8/SVneo cells, but not G9a. G9a was a negative control. (E) Western blot assays detected the expression of *RND3* after silenced *EZH2* after si-RNAs transfection in HTR-8/SVneo cells. (E) The enrichment of *EZH2*/H3K27me3 and *LSD1*/H3K4me2 in the promoter regions of *RND3* were identified via ChIP assays, and this enrichment was decreased after *HOXA11-AS* knockdown in HTR-8/SVneo cell lines. Antibody directed against IgG was used as a negative control. (Values represent the mean **±** S.E.M from three independent experiments. **: P < 0.01, *: P<0.05, n.s.: not significant)

To examine that the interaction probabilities of *HOXA11-AS* with target proteins, we performed RIP assays with these antibodies. There was a substantial enrichment in RIPs of Ezh2, Lsd1 and Ago2 in HTR-8/SVneo (Figure 5C). Additionally, we also conducted RIP assays in JEG3 and JAR cells which were transfected with overexpression plasmid of *HOXA11-AS*. Our results demonstrated that *HOXA11-AS* directly interacted with Ezh2, Lsd1 and Ago2 (Figure 5C). Furthermore, RNA pull down assays further confirmed that *HOXA11-AS* could interacted with Ezh2, Lsd1, and Ago2 in HTR-8/SVneo cells (Figure 5D). These resulting data demonstrated that *HOXA11-AS* could directly bind with Lsd1, Ezh2 and Ago2 in trophoblast cells. Previous studies have reported that Lsd1 and Ezh2 were negative regulators of transcription via the trimethylation of histone 3 lysine 4 (H3K4me2) and the trimethylation of histone 3 lysine 27 (H3K27me3), respectively [33-35]. Therefore, we further explored the mechanism correlation among Ezh2, Lsd1 and *HOXA11-AS* using experimental methods.

We first suppressed the expression of Lsd1 and Ezh2 by effective siRNAs, respectively. The protein level of *RND3* was significantly upregulated after transfected with *EZH2* siRNAs and/or *LSD1* siRNAs in HTR-8/SVneo cells (Figure 5E). Then, we hypothesized that *HOXA11-AS* may recruit Ezh2 and Lsd1 to *RND3* promoter region, resulting in trimethylation of H3K27 and/or H3K4 in this region. Therefore, we performed ChIP assays to detect the enrichment of Ezh2 and H3K27me3, Lsd1 and H3K4me2 in the promoter region of *RND3*. As shown in the figure 5F, the results determined that Lsd1 and Ezh2 protein could be directly recruited to the promoter region of RND3 gene, thus silencing HOXA11-AS. Silencing of *HOXA11-AS further suppressed*Ezh2-mediated H3K27me3 demethylation and Lsd1-mediated H3K4me2 demethylation.

Our previous studies[31] have reported that the expression level of *RND3* was significantly increased in preeclampsic placental tissues compared to that in the controls. In this study, we also found that overexpression of RND3 could inhibit cell proliferation in HTR-8/SVneo and JEG3 cell lines (Figure 6A-6D). Overexpression of *RND3* could also partly reverse *HOXA11-AS*-mediated growth promotion (Figure 6E and 6F). Together, these data suggest that *HOXA11-AS* medicated cell growth could be reversed partly through epigenetic suppression of *RND3* by binding to Ezh2 and Lsd1 in nucleus of trophoblast cells.

**Figure 6:**
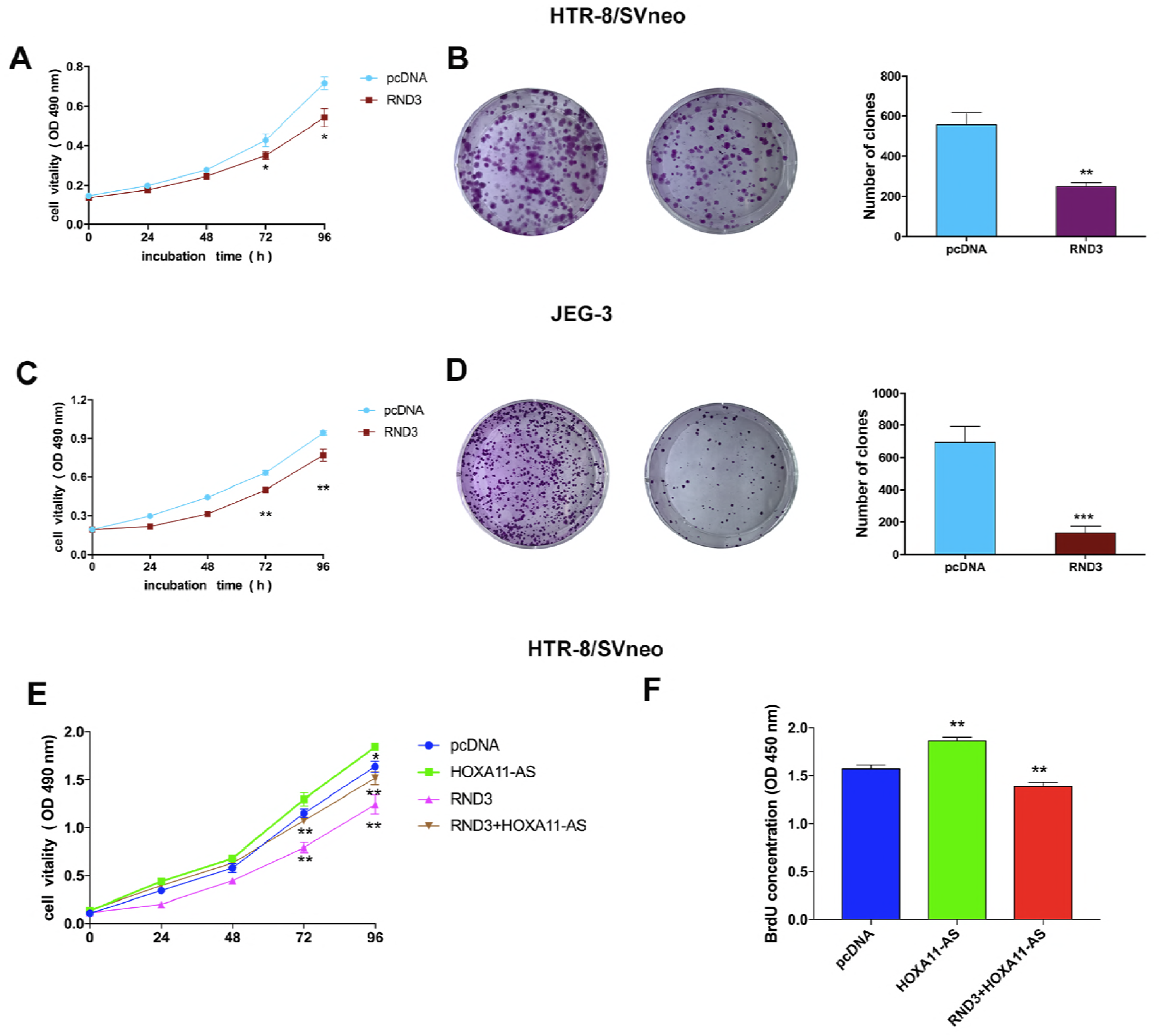
Upregulation of *RND3* inhibit Trophoblast cell proliferation and were involved in the function of *HOXA11-AS*. (A, B, C and D) MTT assays were executed used to determine the cell viability for pcDNA3.1-*RND3*-treated trophoblast cells. Colony-formation assays were performed to assess the cell proliferation for pcDNA3.1-*RND3*-treated HTR-8/SVneo and JEG-3. (E) MTT assays and (F) BrdU assays were implemented to assess the cell viability for RND3 and HOXA11-AS overexpression plasmid co-transfected trophoblast cells. Experiments were performed three times independently. (**: P < 0.01, *: P<0.05)

### HOXA11-AS promotes *HOXA7* expression by sponged miR-15b-5p, thus affecting trophoblast cell proliferation

Based on the RNA-seq analysis, numerous genes affecting cells phenotype were downregulated after silencing of HOXA11-AS. *HOXA7*, part of the cluster on chromosome 7, could promote cell proliferation and migration in various cell lines[36-39]. We next performed Western Blotting assays to further demonstrate the RNA-seq results. As shown in the Figure 7A, we found that the protein level of *HOXA7* was significantly upregulated after *HOXA11-AS* overexpression, whereas the opposite result was found after *HOXA11-AS* knockdown in HTR-8/SVneo cells.

**Figure 7:**
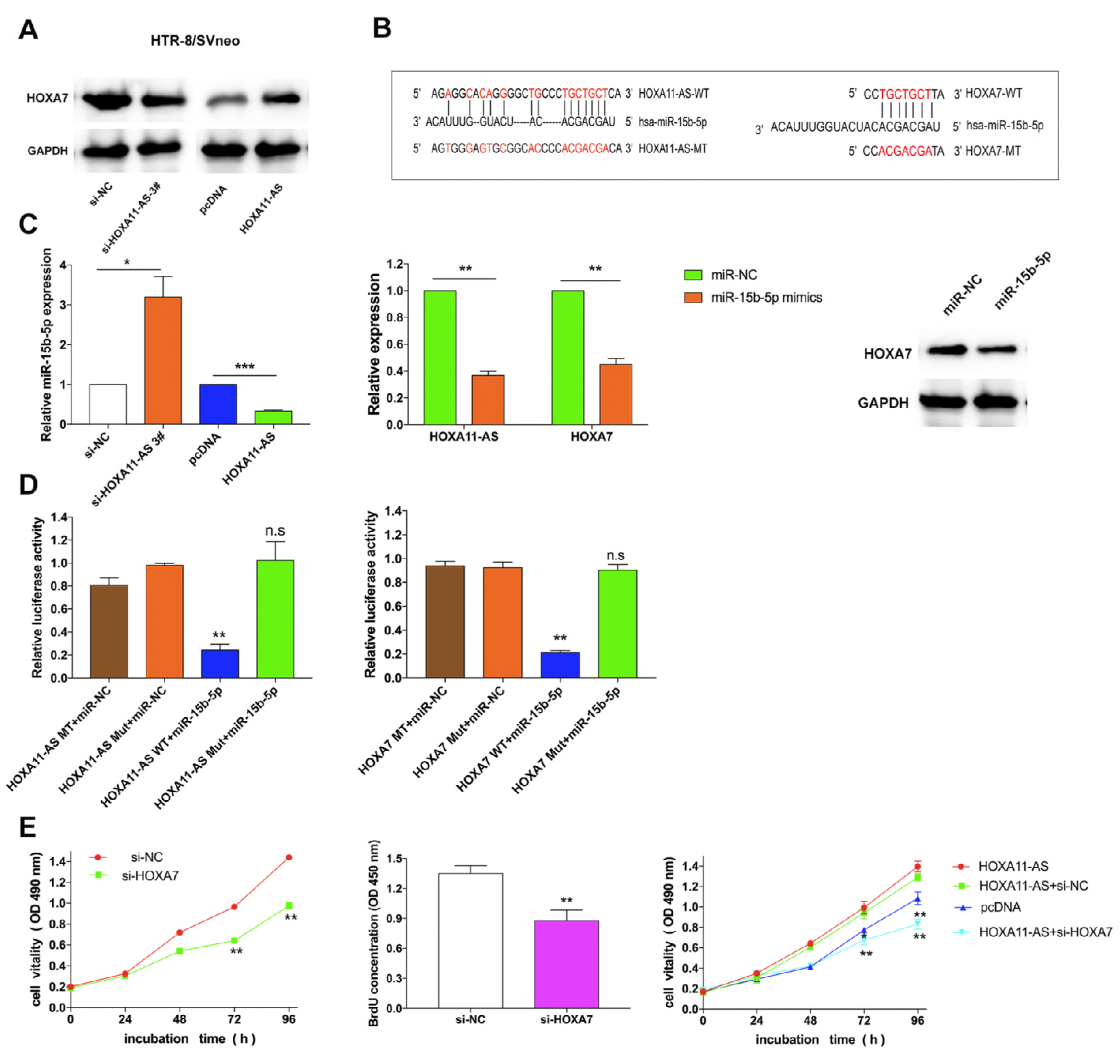
*HOXA11-AS* promotes *HOXA7* expression by competing for miR-15b-5p in cytoplasm, thus facilitating trophoblast cell growth. (A)Western blot assays detected the expression of *HOXA7* after knockdown of *HOXA11-AS* and overexpression of *HOXA11-AS* in HTR-8/SVneo cells. (B) The RNAup algorithm predicted potential binding of miR-15b-5p to *HOXA11-AS* and to *HOXA7*, with considerable sequence complementary in the indicated regions. (C) qRT-PCR assays detected the expression of miR-15b-5p after knockdown of *HOXA11-AS* and overexpression of *HOXA11-AS* in HTR-8/SVneo cells. The expression of *HOXA11-AS/HOXA7* were detected after overexpression of miR-15b-5p in HTR-8/SVneo cells. Western blot assays detected the expression of *HOXA7* after knockdown of overexpression of miR-15b-5p in HTR-8/SVneo cells. (D) Luciferase reporter assays were used to determine the interacting activity between miR-15b-5p and *HOXA11-AS/HOXA7*. Luciferase activity is shown as relative luciferase activity normalized to Renilla activity. (E) MTT and BrdU assays were used to determine the viability of si-HOXA7-transfected HTR-8/SVneo cells, and showed that knockdown *HOXA7* could reverse *HOXA11-AS*-mediated growth. (*P < 0.05, **P < 0.01)

An increasing number of studies have reported that lncRNAs could compete for specific microRNAs (miRNA) in cytoplasm to mediate mRNAs expression, thus further affecting cells phenotype [22]. There is evidence that miRNAs were found predominantly in the cytoplasm by binding to Ago2, which is the fundamental element of the RNA-induced silencing complex. Based on the subcellular fractionation assays and RIP assays (Figure 5C), we found that 30% *HOXA11-AS* were distributed in the cytoplasm and *HOXA11-AS* could interact with Ago2 protein. Then, we hypothesized whether *HOXA11-AS* modulate *HOXA7* expression levels by sponged specific microRNA in HTR-8/SVneo cells.

To validate this concept, we first conducted bioinformatics analysis and found that miR-15b-5p was predicted to bind to the 3′ UTR of downstream target gene of both *HOXA11-AS* and *HOXA7* (Figure 7B). Then, we performed qRT-PCR and the results showed that silenc**ing** of *HOXA11-AS* significantly upregulated the expression of miR-15b-5p; in contrast, the miR-15b-5p level was decreased after *HOXA11-AS* overexpression. Further experiments indicated that RNA and protein levels of *HOXA11-AS* and *HOXA7* were significantly reduced after transfected with miR-15b-5p mimics (Figure 7C). Next, we further explored the functions of miR-15b-5p in trophoblast cells. Diverse luciferase genes, including *HOXA11-AS*, mutant *HOXA11-AS*, 3′ UTR of *HOXA7* and mutant 3′ UTR of *HOXA7*, were cloned and then co-transfected with miR-15b-5p in HTR-8/SVneo cells, respectively. Interestingly, we found that the relative luciferase activity of reporters of *HOXA11-AS* and 3′UTR of *HOXA7* were significantly abolished after treatment by miR-15b-5p (Figure 7D). In contrast, the relative luciferase activity on mutant reporters of both *HOXA11-AS* and 3′UTR of *HOXA7* showed no effect after treatment by miR-15b-5p (Figure 7D). Therefore, these resulting data demonstrated that miR-15b-5p could bind to both *HOXA11-AS* and *HOXA7* gene. Moreover, MTT and BrdU assays indicated that knockdown of *HOXA7* could inhibit proliferation in HTR-8/SVneo cells, and silencing of *HOXA7* could reverse *HOXA11-AS*-induced cell proliferation (Figure 7E).

## Discussion

Recent studies have indicated that plenty of lncRNAs have critical roles in PE. For example, the proverbial lncRNA *MALAT1* were recently determined to affect trophoblast cells proliferation, apoptosis, thus promoting PE development[16]. Our previous study also demonstrated that lncRNA *SPRY4-IT1* and *MEG3* modulate trophoblast cell proliferation, apoptosis as well as tube formation, invasion and migration in this disorder [14, 40, 41]. Moreover, other lncRNAs, including *ATB*, Uc.187, *RPAIN* and so on, were reported to regulate trophoblast cell growth or/and invasion or/and apoptosis in PE[17, 18, 42]. Therefore, more and more potential PE-associated lncRNAs need to be identified, the functions of which require further exploration for their underlying biological mechanisms.

In our study, we discovered that the level of *HOXA11-AS* was significantly downregulated in preeclampsic tissues compared with those in normal tissue samples, thus suggesting that *HOXA11-AS* might play an essential role in PE progression. Our data also revealed that knockdown of *HOXA11-AS* could impair trophoblast cell proliferation and migration *in vitro*, whereas *HOXA11-AS* overexpression could promote cell proliferation and migration. In order to investigate the *HOXA11-AS* -related pathway and downstream genes in PE, we conducted RNA transcriptome sequencing after transfecting target cell line with specific siRNAs against *HOXA11-AS*, and GO analysis suggested that gene expression profiles were primarily proliferation and migration associated. Previous studies have determined that lots of lncRNAs could bind with various chromatin modifying enzymes to regulate related genes expression at epigenetic level [43]. For instance, the pseudogene *DUXAP10* promotes an aggressive phenotype through binding with *LSD1* to repress *LATS2* and *RRAD* in non-small cell lung cancer [44], and LncRNA *TUG1* was involved in cell proliferation of small cell lung cancer by regulating *LIMK2b* via *EZH2* [45]. Our resulting data also revealed that *HOXA11-AS* could recruit and bind to two histone methylation modification complexes, including *EZH2* and *LSD1* in nucleus, thus silencing genes expression. Further experiments were performed, indicating that these targets were affected through promoter H3K4me2 demethylation and H3K27me3 demethylation in trophoblast cells.

*RND3/ RhoE* is a small GTPase that could exhibit biological functions as suppressor genes in numerous diseases[46-49], thus inhibiting multiple cellular processes, such as actin cytoskeleton dynamics, cell cycle[46], migration, invasion[50], and apoptosis[46, 51]. However, the functions of RND3 in the pathological process of PE was still unclear. Our results revealed that *HOXA11-AS* could contribute to the downregulated expression of *RND3* by histone methylation in trophoblast cells. These results suggested that *HOXA11-AS* could bind to Ezh2 and Lsd1, thus epigenetically silencing *RND3* in nucleus of trophoblast cells.

HOX genes, a highly conserved subgroup of the homeobox superfamily, are spatially and temporally regulated during embryonic development[52]. Abnormal expression of *HOXB7*[53] and H*OXB13* [54] have been reported to regulate many processes including apoptosis, receptor signaling and differentiation in myriad of disorders[55]. In our study, we found that low levels of *HOXA7* could affect proliferation in trophoblast cell lines, which implies that the post-transcriptional regulation of *HOXA7* was partly mediated by *HOXA11-AS* in development of PE, through sponging miR-15b-5p in cytoplasm, thus accelerating trophoblast cell growth. Furthermore, Sun et al [29] has demonstrated that *HOXA11-AS* could promote gastric cancer tumorigenesis through sponging miR-1297. These findings demonstrated that *HOXA11-AS* simultaneously compete for miR-1297 and miR-15b-5p.

In brief, *HOXA11-AS* facilitates cell proliferation and migration by epigenetically regulating RND3 in nucleus. And it could promote trophoblast cell growth through sponging miR-15b-5p in cytoplasm. Our results indicated that *HOXA11-AS* could exhibit diverse biological regulatory mechanisms in PE, which might act as a prospective diagnosis and therapeutic target for PE (Figure 8).

**Figure 8.**
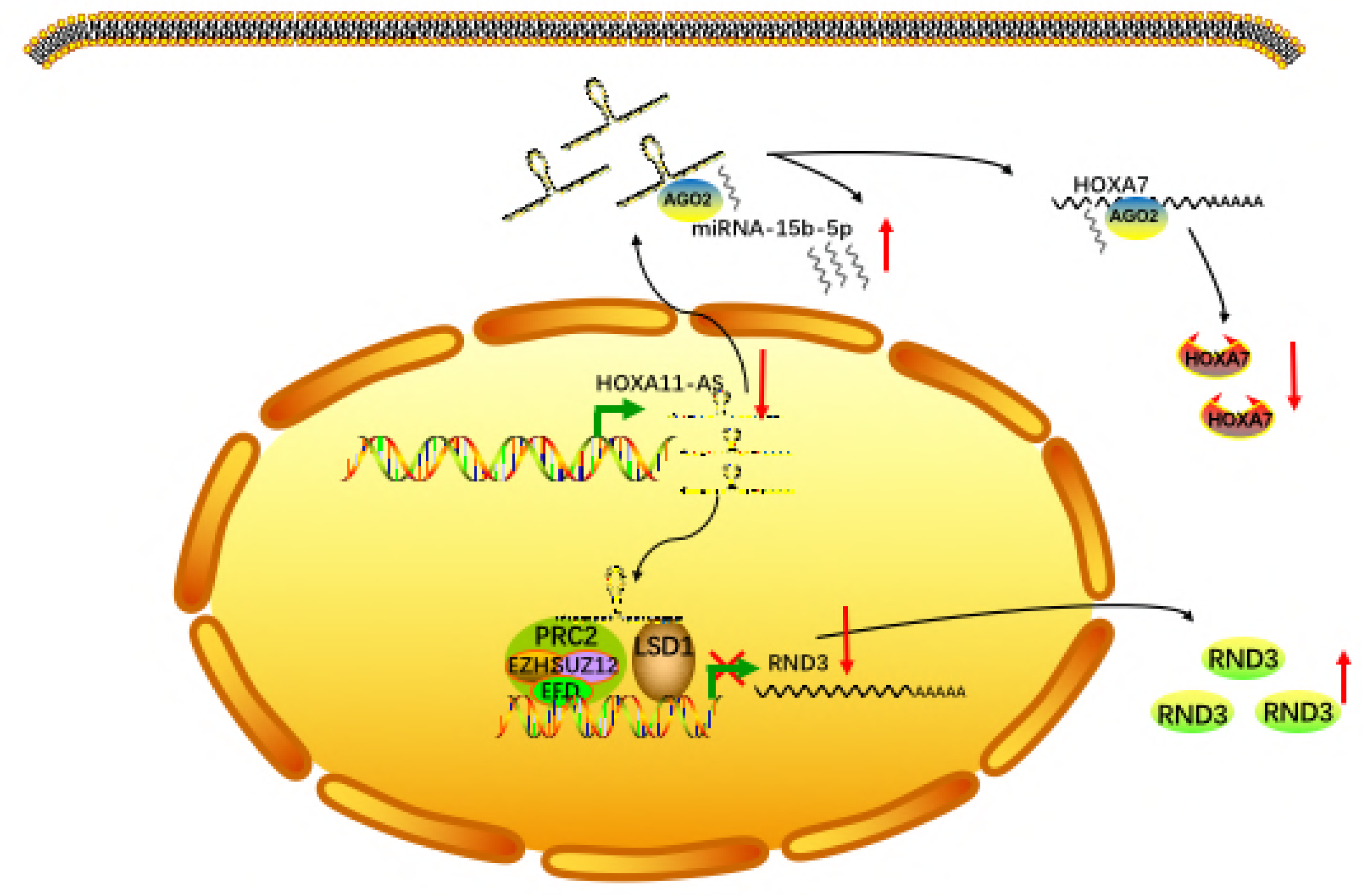
Proposed model which medicated by HOXA11-AS in proliferation and migration progression of PE.

## Additional file

Table S1. Sequence of primers and siRNAs.

Table S2 and S3. Analysis of the RNA transcriptome sequencing data.

## Author Contributions

Yetao Xu, Dan Wu and Jie Liu performed the most experiments. Shiyun Huang, Qing Zuo and Yifan Lian collected clinic tissues, analyzed data. Zhonghua Ma, Bingqing Hui, Yanzi Chen and Tianjun Wang conducted some experiments. Yetao Xu and Lizhou Sun designed the project and edited the manuscript.

## Abbreviations

lncRNA: (long non-coding RNA)
miRNA: (micro RNA)
PE: (Preeclampsia)
EZH2: (enhancer of zeste homolog 2)
LSD1: (Histone demethylase lysine specific demethylase 1)
Ago2: (argonaute 2)
RND3: (Rho family GTPase 3)

## Acknowledgments

This study was supported by the National Scientific Foundation of China (No. 81270710, NO. 81470065 and NO.81771603), the traditional Chinese medicine project of Jiangsu Province (NO. ZX2016D2) and the project of construction capacity for birth defect screening and diagnosis laboratory in Jiangsu (BM2015020), and the Natural Science Foundation of Jiangsu Province (project number: BK20161061 and BK20171502).

